# Distinct core minor pilin complexes prime specialized type IV filaments in cyanobacteria

**DOI:** 10.64898/2026.02.12.705559

**Authors:** Nils Schuergers, Lars Wittig, Gen Enomoto, Julia Herz, Shamphavi Sivabalasarma, Katharina Ruoff, Friedel Drepper, Sonja-Verena Albers, Pitter F. Huesgen, Annegret Wilde

**Affiliations:** Molecular Genetics of Prokaryotes, Faculty of Biology, University of Freiburg, 79104 Freiburg, Germany; Department of Agricultural Chemistry, Tokyo University of Agriculture, Tokyo, 156-8502 Japan; Molecular Biology of Archaea, Faculty of Biology, University of Freiburg, 79104 Freiburg, Germany; CIBSS-Centre for Integrative Biological Signaling Studies, University of Freiburg, 79104 Freiburg, Germany; Biochemistry and Functional Proteomics, Faculty of Biology, University of Freiburg, 79104 Freiburg, Germany

## Abstract

The model cyanobacterium, *Synechocystis* sp. PCC 6803 encodes in addition to the major pilin of the Type IV pilus filament, an extensive, partially uncharacterized repertoire of minor pilins. For those cyanobacterial minor pilins that have been characterized, their roles span a surprisingly diverse range of functions. To elucidate the roles of uncharacterized minor pilins in a systematic way, we applied structural phylogenomics across 90 genomes, classifying cyanobacterial pilins into six conserved families that form two distinct putative core priming complexes. We demonstrate that these complexes initiate the assembly of two morphologically distinct Type IV pilus filaments: short, hyper-dynamic pili for natural competence, and long, adhesive pili for motility and phototaxis. Structural analysis revealed a conserved beta-solenoid domain in PilX subunits, which we propose fulfills the “tip plug” function of PilY1 homologs. Proteomic data indicate that the minor pilin PilX2 facilitates the assembly of motility pili, which is required to maintain DnaJ3 co-chaperone levels and trigger cAMP-dependent surface sensing. These findings challenge the concept of a single multipurpose pilus, establishing that cyanobacteria operate two specialized nanomachines optimized for the conflicting biophysical requirements of DNA uptake and surface motility.

## Introduction

The Type IV filament superfamily comprises ubiquitous nanomachines, including Type IV pili and Type II secretion systems (T2SS), found throughout bacteria and archaea (1, 2). The most widespread subfamily, Type IVa pili (T4P), are dynamic appendages that undergo rapid cycles of extension (polymerization) and retraction (depolymerization), driven by a membrane-spanning T4P-machinery (3–5). This activity enables diverse processes, including surface-dependent translocation, known as twitching motility, adhesion, biofilm formation, and uptake of exogenous DNA during natural competence (6, 7).

The T4P filament is a helical polymer composed primarily of thousands of copies of a single major pilin that forms the structural backbone of the pilus (8–10). It also incorporates structurally diverse, low-abundance minor pilins (11). All pilins are synthesized as prepilins with an N-terminal type III signal peptide that is cleaved by the prepilin peptidase PilD (12). Mature pilins adopt a characteristic ‘lollipop’ fold, in which a conserved hydrophobic N⍰terminal α⍰helix (α1N) anchors the prepilin in the inner membrane and later forms the hydrophobic core of the assembled fiber. The C⍰terminal portion of the helix contributes to the globular domain, whose base fold is an antiparallel β⍰sheet extended by variable regions (13, 14).

In heterotrophic model organisms, a set of “core” minor pilins (FimU/PilV/PilW/PilX in *Pseudomonas aeruginosa*) typically forms a stoichiometric tip complex —also known as the priming complex—because it primes polymerization of the T4P filament (15–21). Structural homology with the T2SS implies a tip-to-base assembly logic: PilV and PilW interact first, followed by the distal subunit PilX, before the adapter FimU anchors the complex to the main fiber (17, 20, 22, 23). Critically, the terminal subunit PilX lacks the conserved Glu5 residue required for subunit addition, effectively capping the filament (8, 13). In Proteobacteria, tip-associated adhesins, such as PilY1 or PilC, are proposed to act as terminal “plugs,” and mechanosensors that couple surface engagement to downstream signaling (18, 20, 24–27).

Additional pilins that are dispensable for filament assembly but are critical for conferring specific functionalities are termed non-core minor pilins (11). These accessory pilins adapt T4P for diverse roles, such as facilitating adhesion or acting as specific receptors for DNA uptake in naturally competent bacteria. While some are predicted to be part of the pilus tip, others are thought to be incorporated throughout the pilus filament (15, 28–30).

While the functional repertoire of minor pilins is increasingly appreciated in heterotrophic model organisms, the cyanobacterial machinery remains poorly defined (31). In the unicellular photosynthetic model *Synechocystis* sp. PCC 6803 (hereafter *Synechocystis*), T4P drive phototactic twitching motility, cell-cell adhesion (flocculation), and natural competence (32–34). This functional complexity is reflected in a genome encoding the major pilin PilA1, along with at least 13 additional putative pilins. Initial characterization suggests functional specialization: PilA5 is essential for transformation but dispensable for motility, whereas the locus encoding PilA9-12 is critical for motility but irrelevant for competence (34–36). Evidence of pilin function in other cyanobacteria remains scarce, mainly limited to the description of pilin-like proteins that facilitate natural transformation in *Synechococcus elongatus* (37) or motility of hormogonia, specialized filaments of *Nostoc punctiforme* (38).

This raises the question of whether cyanobacteria possess a single T4P machine incorporating specialized accessory pilins or assemble multiple, distinct filaments. Addressing this has been hindered by the extreme sequence divergence of minor pilins. Here, we combined structural phylogenomics with functional genetics to demonstrate that *Synechocystis* utilizes distinct sets of core minor pilins to prime two morphologically distinct filaments, thereby distinguishing the machinery required for motility from that involved in DNA uptake.

## Results

### Characterization of minor pilin mutant strains

While the *Synechocystis* minor pilins PilA2, PilA5-6, PilA9-12, PilX1, PilX2, and PilX3 have been characterized previously (33–36), the roles of PilA7-8 and PilA4 remain unknown. To obtain a full picture of minor pilin functions, we constructed knockout mutants and complementation strains for these genes in the motile *Synechocystis* PCC-M background strain (39). Concurrently, a Δ*pilA2* mutant was generated to investigate its role in cellular aggregation and verify previously established characteristics in another wild-type background (32, 33).

Deletion of the *pilA7-8* transcriptional unit (Δ*pilA7-8*) revealed no discernible effect on phototactic motility; the mutant moved toward light, similar to the wild type (WT) (Fig. 1A). This phenotype persisted in complementation strains expressing PilA7, PilA8, or both in *trans* (Δ*pilA7-8*/*pilA7*^*+*^, Δ*pilA7-8*/*pilA8*^*+*^, and Δ*pilA7-8*/*pilA7*-*8*^*+*^, respectively). These findings are consistent with electron microscopy observations confirming that the Δ*pilA7-8* mutant retained surface pili (Fig. S1). However, the Δ*pilA7-8* strain exhibited significant hyper-aggregation compared with the WT, whereas Δ*pilA1* mutant cells did not aggregate (Fig. 1B). Notably, the lack of aggregation in our Δ*pilA1* mutant contrasts with previous reports, yet is fully consistent with the dependence of flocculation on pilus assembly by the PilB1 extension motor (34).

**Figure 1.**
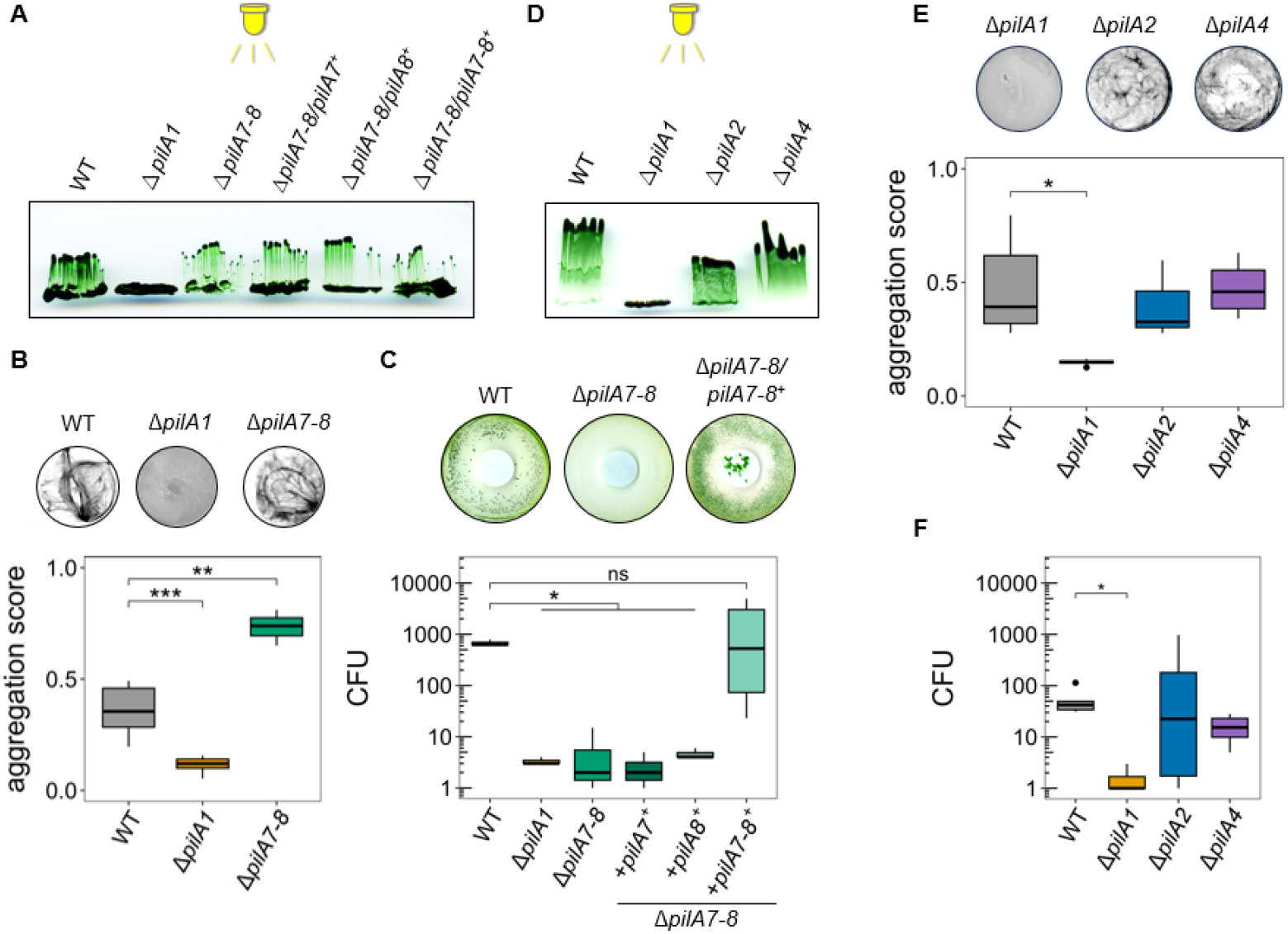
Phenotypic characterization of minor pilin mutants reveals that PilA7 and PilA8 are crucial for natural competence but not for motility. **(A, D)** *Synechocystis* strains were restreaked on 0.5% (w/v) BG-11 motility agar and exposed to directional white light placed in directional white light (∼ 15 µmol photons m^−2^ s^−1^, indicated by the LED symbol) to assess phototactic motility. **(B, E)** Cultures in 6-well plates were incubated for 48 h with mild orbital shaking, and chlorophyll autofluorescence was imaged (upper panel). The aggregation score of at least three replicates was calculated by dividing the standard deviation of chlorophyll autofluorescence intensity by the mean intensity (lower panel). **(C, F)** Natural transformation efficiency was measured by counting the number of antibiotic-resistant colonies (CFU) appearing in the inhibition zone two weeks after transformation with 500 ng of a plasmid (upper panel). CFU were counted for three replicates. Statistical significance between WT and mutant strains was assessed using pairwise t-tests (p-values < 0.05*, <0.01**, <0.001***).

Although motility and aggregation remained intact, the Δ*pilA7-8* mutant was found to be no longer naturally transformable (Fig. 1C). This deficiency could not be rescued by the individual expression of PilA7 or PilA8. Only the expression of both PilA7 and PilA8 in Δ*pilA7-8*/*pilA7-8*^*+*^ successfully restored transformation efficiency to WT levels, demonstrating that PilA7 and PilA8 are collectively essential for natural competence. This functional profile resembles that of the upstream-encoded minor pilin PilA5, suggesting that the *pilA5-6* and *pilA7-8* transcriptional units together form a dedicated DNA uptake locus.

Finally, the Δ*pilA4* mutant showed no defects in piliation, phototactic motility, aggregation, or DNA uptake (Fig. 1D–F and Fig. S1). The Δ*pilA2* strain displayed normal phototaxis consistent with previous reports (32, 33), whereas its transformation efficiency varied between clones and was not consistently reduced. Furthermore, we observed that the absence of *pilA2* had no discernible effect on aggregation (Fig. 1E).

### Structural clustering reveals two conserved minor pilin sets in cyanobacteria

To resolve the evolutionary relationships of cyanobacterial pilin proteins, we compared the predicted structures of putative pilins from a diverse set of 90 cyanobacterial species. We applied the structural alignment and hierarchical clustering workflow (Fig. 2A) detailed in the Materials and Methods section, yielding 13 statistically supported clusters (AU p-value > 0.95) that we could consolidate into six major families: the major pilin PilA family, a related auxiliary pilin family, and families with homology to the four core minor pilins, that is, the FimT, PilW, PilV, and PilX families (Fig. 2B, Datasets S1 & S2).

**Figure 2.**
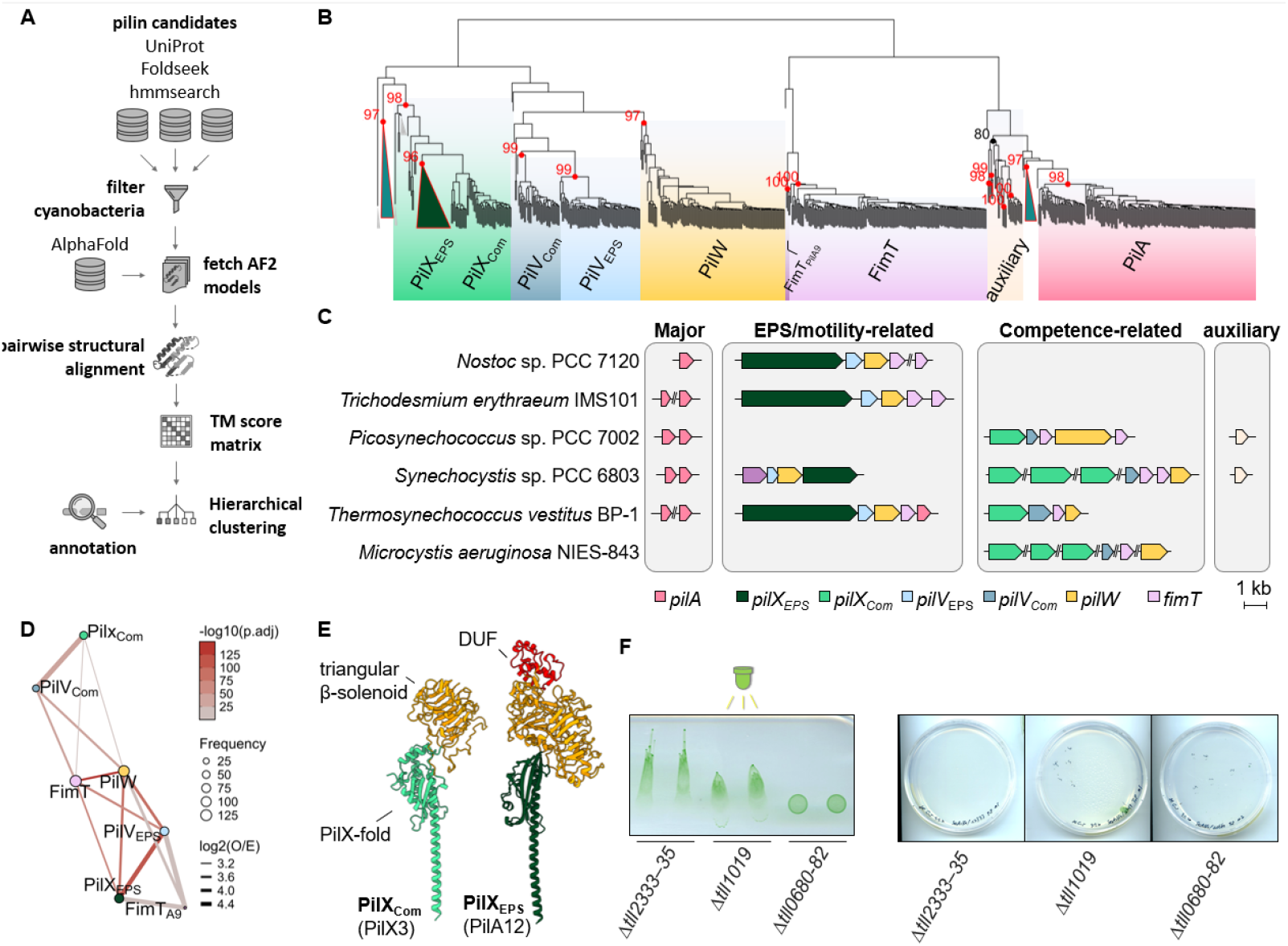
Structural clustering classifies cyanobacterial pilins into six families and identifies conserved sets that drive motility and competence. **(A)** Computational workflow for classifying cyanobacterial pilins. Putative pilin sequences were aggregated using annotation-, homology-, and structure-based searches (UniProt, HMMER, Foldseek). After filtering to retain reference genomes, 792 AlphaFold2-modeled candidate structures were aligned pairwise using USalign. The resulting similarity matrix was subjected to hierarchical clustering (pvclust) to define statistically supported pilin families. **(B)** Hierarchical clustering dendrogram. Node labels represent approximately unbiased (AU) p-values (multiscale bootstrap). Shading indicates pilin families, triangles mark collapsed nodes: PilX_EPS_ clade containing unmodeled homologs (dark green), pilin-like proteins missing the α1N helix (cyan), and non-pilin proteins (grey). **(C)** Genomic organization of major and minor pilins, highlighting the distribution of two conserved core minor pilin sets across representative cyanobacteria. Scale breaks indicate that the displayed genes are not adjacent in the genome (see Datasets S1 & S2 for details). **(D)** Network showing co⍰occurring pilin families encoded together in the same gene clusters. Nodes are scaled by frequency of observation, while connecting edges denote significant positive associations (Fisher’s Exact Test, BH-adjusted p < 0.05, log_2_(Obs/Exp) > 1). **(E)** AlphaFold predictions of PilA12 (AF⍰P73238) and PilX3 (AF⍰P73549). The core pilin fold is shown in green (distinct shades for each subfamily), the bulky apical β⍰solenoid domain in orange, and additional domains of unknown function (DUF) in red. **(F)** Functional analysis of *Thermosynechococcus* minor pilin deletion mutants. (Left) Phototactic motility was assayed on soft agar plates exposed to directional green light of 120 µmol photons m^-2^ s^-1^ for 24 h. (Right) Natural transformation efficiency was assessed by colony recovery on kanamycin-supplemented agar (80 µg ml-1) after incubation with plasmid DNA.

The PilA family constitutes the canonical major pilin cluster. A divergent family of ‘auxiliary pilins’ was also identified, though with weak statistical support. We chose this specific nomenclature because these proteins are likely unnecessary for assembly—as seen with PilA4—despite their structural similarity to the major pilin. The FimT family split clearly into a major cluster and a minor group containing *Synechocystis* PilA9. Crucially, the clustering analysis revealed a deep split within the PilV and PilX families, which clearly defined two subfamilies. Consistent with this divergence, a strict co-occurrence pattern was observed between the PilX and PilV subfamilies, even when the genes were not colocalized, suggesting a functional linkage (Fig. 2C, D). All cyanobacterial PilX homologs feature a canonical pilin fold interrupted by a bulky, triangular β-solenoid domain that resembles ice-binding proteins and other adhesive domains inserted between the penultimate and C-terminal β-strands (Fig. 2E). In one subfamily, this domain is significantly expanded and homologous to HpsA, a protein essential for motility and hormogonium polysaccharide (HPS) secretion in *Nostoc punctiforme* hormogonia (40). Although only a few cyanobacterial strains differentiate hormogonia, extracellular polysaccharides (EPS) are likely universally required for T4P-mediated motility in cyanobacteria. Consequently, we designated this subfamily as PilX_EPS_ and the co-occurring PilV clade as PilV_EPS_. The remaining PilX variants are associated with the PilV subfamily that includes competence-associated pilins such as *Synechocystis* PilA5 and *S. elongatus* RntB. Consequently, we designated these subfamilies as PilX_Com_ and PilV_Com_.

Our analysis identified an additional class of pilin-like proteins, distributed across two separate clades, that lack the conserved α1N-helix. These proteins possess accessory domains, such as calycin-type β-barrels or serine/threonine kinase domains, suggesting that they may function as regulators rather than structural subunits of the pilus filament (Datasets S1 & S2 and Fig. S2).

### Two conserved minor pilin sets are encoded in cyanobacterial genomes

A critical insight from the clustering analysis was the strict genomic co-occurrence of PilX_EPS_ with PilV_EPS_ and of PilX_Com_ with PilV_Com_, suggesting that these proteins function as obligate structural pairs, likely forming distinct priming complexes that serve as nucleation seeds for their respective Com-, or EPS-type filaments.

Genomic analysis across cyanobacterial strains has revealed substantial variability in pilin gene numbers, ranging from none or one in some species to as many as 44 in *Acaryochloris marina*. Most genomes, however, encode between 7 and 11 putative pilins (Fig. 2C and Datasets S1 & S2). Strains typically possess two major pilin genes, regularly located adjacent to a gene encoding an O-linked N-acetylglucosaminyltransferase (*ogt*) crucial for pilin glycosylation (41, 42), and complete sets of core minor pilin genes (PilX-PilV-PilW-FimT) often encoded within a single locus. While some species encode only a single Com- or EPS-type set of pilins, others encode both, and some, such as *A. marina*, encode multiple sets. In addition, some species encode, in most cases, a single auxiliary pilin.

To experimentally validate the functional link between these conserved sets and natural competence or cell motility, we constructed pilin deletion mutants in *Thermosynechococcus vulcanus*. Targeting the predicted PilX_EPS_-PilV_EPS_ encoding EPS/motility-related locus (*tlr0680-82*) abolished phototactic movement, whereas targeting the predicted PilX_Com_-PilV_Com_ encoding competence-related locus (*tll2333-35*) resulted in a complete loss of natural transformability, with no discernible effect on motility (Fig. 2F). Deletion of the second major pilin homolog, *tll1019*, caused a reduction in motility. These results confirm that the functional specialization of minor pilins observed in *Synechocystis* is conserved in *T. vulcanus*.

While our structural classification clearly delineates two distinct lineages, the designations “EPS” and “Com” reflects the evolutionary divergence of these clusters rather than a strict functional separation. Genomic and experimental evidence suggest a degree of functional plasticity or cross-talk between these systems. For instance, although *Synechocystis* PilX2 belongs to the Com-type lineage, its deletion results in a discernible motility defect (36). Furthermore, the robust motility of *Synechococcus elongatus* UTEX 3055—which lacks the EPS-associated cluster and relies solely on two Com-type minor pilin sets (43)—indicates that the function of the core minor pilins may vary among species.

### The two core minor pilin sets of *Synechocystis* can independently prime pilus assembly

The functional differentiation and persistence of surface pili in single-operon mutants suggest that the two core minor pilin sets prime separate pilus types. To test whether these two sets independently support pilus assembly, we analyzed the presence of surface pili in various mutant strains. Therefore, the presence of PilA1 in surface-sheared protein samples was compared to intracellular PilA1 levels to rule out defects in pilin expression (Fig. 3A). Consistent with our earlier findings, the Δ*pilA9-12* (lacking the EPS set) and Δ*pilA5-6* and Δ*pilA7-8* mutants (lacking components of the Com set) all retained detectable PilA1 in the sheared surface protein fraction, with the Δ*pilA2* mutant showing a statistically significant increase in extracellular PilA1 levels (adj. P = 0.00053). In sharp contrast, no surface PilA1 signal was detected in protein fractions sheared from either the Δ*pilA5-6*/Δ*pilA9-12* or the Δ*pilA7-8*/Δ*pilA9-12* double mutants. As no significant variation in intracellular PilA1 abundance was observed, these differences demonstrate that a single functional set of core minor pilins, either Com or EPS, is sufficient to prime pilus assembly.

**Figure 3.**
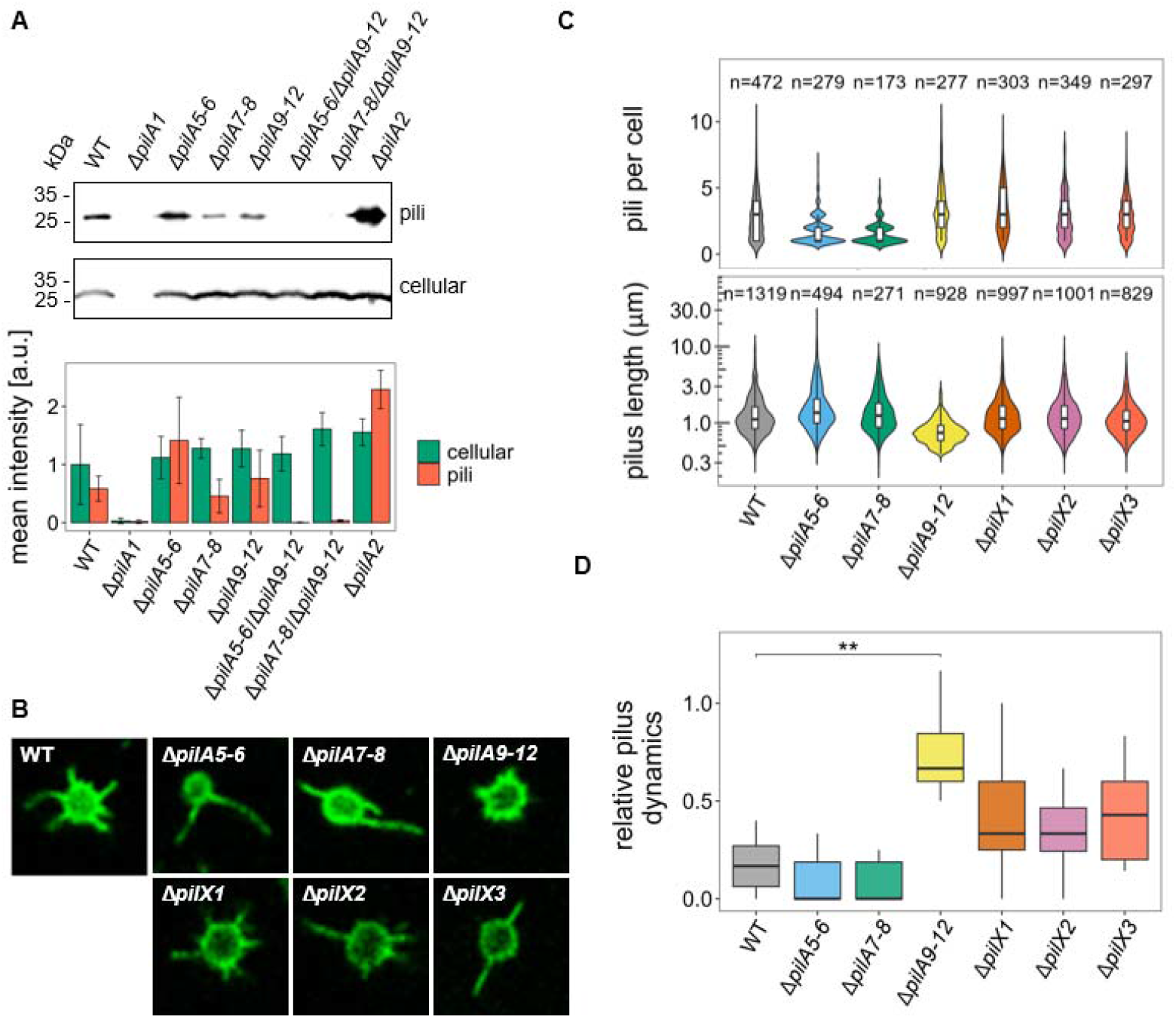
Core minor pilin sets independently assemble morphologically distinct pilus filaments. **(A)** Representative western blot analysis of PilA1 abundance in sheared surface fractions and whole-cell lysates (top) and signal quantification from three independent experiments with error bars representing standard deviation (bottom). Surface pili were isolated from the indicated *Synechocystis* mutant strains, separated by 15% SDS-PAGE, and probed with an anti-PilA1 antibody. The Δ*pilA1* mutant was used as a negative control. Surface PilA1 was absent only in double mutants. **(B)** Representative confocal microscopy images of live cells visualized on 2% agar pads. All images were acquired using the PilA1(T77C) background required for thiol⍰specific maleimide labeling, and pili were labeled with Alexa Fluor 488 maleimide for 2 h. **(C)** Quantification of pilus morphology. Semi-automated image analysis was used to determine the number of pili per cell (upper panel) and pilus length (lower panel). Mutants retaining only the PilA9-12 set (Δ*pilA5-6* and Δ*pilA7-8*) displayed fewer but longer pili, whereas the mutant retaining only the PilA5-8 set (Δ*pilA9-12*) exhibited a significantly higher number of short pili than the wild type (WT). **(D)** Quantification of pilus dynamics. The relative change of piliation was calculated for 10 cells as the sum of assembly and disassembly events over 20 s, normalized to the initial pilus count per cell. The short pili assembled by the PilA5-8 set (Δ*pilA9-12*) were significantly more dynamic compared to the WT (T77C-Strep) (pairwise Wilcoxon rank sum test, ** p < 0.05).

### EPS and Com minor pilins prime morphological distinct filaments

To visualize potential differences in pilus morphology and dynamics *in vivo*, we employed a staining strategy utilizing thiol-reactive cysteine variants of the major pilin PilA1 (Fig. S3). Fluorescence microscopy of strains expressing a PilA1(T77C) variant that retained pilus dynamics in different minor pilin mutants revealed striking morphological differences that were quantified by semi-automated image analysis (Fig. 3 B, C). Mutants assembling pili exclusively via the EPS set (i.e. Δ*pilA5-6* or Δ*pilA7-8*) produced fewer, longer pili than the WT. Conversely, the Δ*pilA9-12* mutant, which assembles pili exclusively via the Com set, displayed a significantly higher number of substantially shorter pili. In contrast, differences in the *pilX* mutants were less pronounced, although the Δ*pilX1* mutant showed a statistically significant increase in pilus number, and the Δ*pilX3* mutant exhibited increased pilus length (Fig. 3 B, C).

Next, we analyzed pilus dynamics, defined as the total change in pilus number within a 10-second interval (see example Movies S1-7). This quantification (Fig. 3D) showed that the short Com filaments were significantly more dynamic than the longer EPS filaments observed in the other strains. Since the formation of new pili was rarely observed, the differences in dynamics were primarily attributed to variations in pilus retraction. This suggests a facilitated retraction mechanism for the short, highly dynamic Com-pili, possibly due to the absence of surface attachment. On the other hand, the longer dwelling times of EPS pili are consistent with a function in surface attachment and motility.

### PilX2 links motility pilus assembly to cAMP and the DnaK1-DnaJ3 system

We performed quantitative proteomics on sheared pilus fractions to dissect the composition of pilus subtypes and identify the PilX paralogs that cap the Com-type tip complex (Fig. 4 & Dataset S3). We reasoned that the deletion of specific minor pilins would prevent the assembly and subsequent secretion of their respective tip complexes, leading to the depletion of their constituent subunits from the extracellular fraction. However, differential abundance analysis revealed that PilX profiles did not exhibit simple depletion patterns. While PilX1 and PilX3 were depleted in all minor pilin mutants, PilX2 showed only marginal increases in some cases. These overlapping fluctuations precluded assigning PilX variants to defined complexes based solely on abundance in sheared fractions.

**Figure 4:**
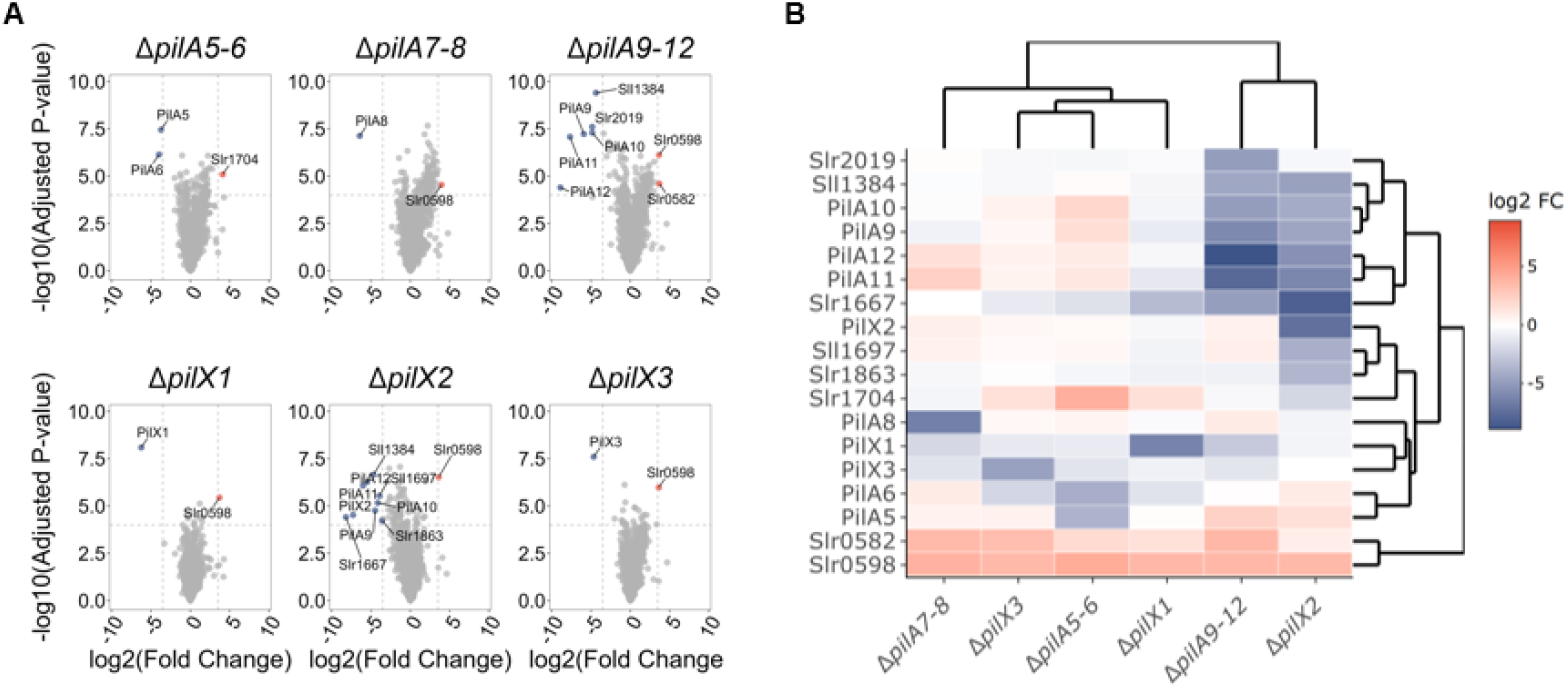
Quantitative proteomic profiling of the sheared T4P fraction in *Synechocystis* minor pilin mutants. **(A)** Combined volcano plots comparing the proteomic composition of the extracellular sheared protein fraction between the indicated mutants and the WT (n=4; n= 3 for Δ*pilA7-8*). Cells were grown on glucose-supplemented plates for 8 days to promote piliation, resuspended in PBS, and subjected to mechanical shearing. The resulting supernatants were precipitated using ammonium sulfate, resolubilized in 2% SDS, and processed via SP3-based tryptic digestion prior to LC-MS/MS. The x-axis represents the log_2_ fold change (FC) relative to the WT. Red indicates significant enrichment (log_2_ FC > 3.5, adj. *p* < 10^-4^) and blue indicates significant depletion (log_2_ FC < -3.5, adj. *p* < 10^-4^. **(B)** Hierarchical clustering heatmap of proteins with significant variance across all comparisons. The color scale represents the log_2_ FC relative to the WT, ranging from depletion (blue) to enrichment (red).

Nonetheless, we observed a reciprocal accumulation pattern: enrichment of Com pilins (PilA5-8) coincided with the depletion of EPS pilins (PilA9-12), suggesting that the cell allocates a fixed pool of the membrane-spanning T4P-machinery between the two filament types. Cluster analysis grouped Δ*pilX1* and Δ*pilX3* with Δ*pilA5-6* and Δ*pilA7-8* mutants, supporting their association with the Com pilus. However, the profile of the sheared pili fraction of Δ*pilX2* clustered with that of Δ*pilA9-12*. Specifically, Δ*pilX2* sheared fractions were significantly depleted of EPS pilins (PilA9-12) (Fig. 4B). Furthermore, the high sensitivity of the MS analysis resulted in the co-detection of intracellular proteins (likely released via cell lysis), allowing us to observe some broader proteomic changes. Both Δ*pilX2* and Δ*pilA9-12* mutants showed downregulation of proteins controlled by cAMP (e.g., Slr1667) and the chaperone DnaJ3 (Sll1384), which is essential for motility but not pilus assembly (44, 45). Since DnaJ3 is not part of the known cAMP regulon, its downregulation likely reflects a systemic feedback mechanism triggered by the loss of the motility pilus. This implies that despite its structural grouping with the Com-set, PilX2 is important for the biogenesis of the motility pilus.

## Discussion

The function of the extensive pilin repertoire in cyanobacteria remains an open question, obscured by the sequence divergence of these proteins. By classifying pilin structures across 90 cyanobacterial genomes, we showed that they fall into just six families. The supramolecular priming complex (FimT–PilW–PilV–PilX) constitutes a universal architecture, with functional specializations (Com vs. EPS) layered upon it. This challenges the notion that *Synechocystis* relies on a single adaptable T4P filament, functionally tailored for diverse tasks through the incorporation of accessory pilins. Instead, our data support a “distinct pilus” model, in which the T4P machinery assembles two distinct filaments. This structural framework allows for the interpretation of pilin-dependent phenotypes in other cyanobacteria. For instance, in *S. elongatus*, proteins PilA3, PilW, RntA, and RntB—previously identified as essential for natural transformation (37)—can now be annotated as a complete Com-type priming complex (see Datasets S1 & S2), whereas a second minor pilin set governs biofilm-associated behaviors (46).

The assembly of distinct filaments allows the bacterium to decouple their biophysical properties, allowing the optimization for disparate functions. The Com filament primed by PilA5-8 is short, abundant, and hyper-dynamic. We hypothesize that this morphology is biophysically optimized for DNA scavenging, where rapid extension and retraction cycles increase the sampling frequency required to capture DNA molecules. Conversely, the EPS filament (primed by PilA9-12) is long, sparse, and persistently attached. This architecture supports a grappling hook function, enabling the pili to bridge distances and sustain the tension necessary for motility or aggregation. These observations mirror the functional partitioning observed in other phyla. For instance, *Thermus thermophilus* utilizes a single T4P machine to assemble wide and narrow filaments from distinct pilins (47), alongside putative short DNA-uptake pseudopili (48), while *Streptococcus sanguinis* generates both straight/thin and wavy/thick filaments from the same major pilin pool (49). Together, these observations are consistent with an evolutionary strategy in which filament architecture is optimized for specific physiological functions.

An interesting finding is that cyanobacterial PilX homologs diverge from the smaller PilX homologs of Proteobacteria by incorporating a bulky β-solenoid domain into the canonical pilin fold – an architecture also found in the tip-associated pilin ComZ of *Thermus thermophilus* (18). In heterotrophic models, the tip adhesin PilY1/PilC serves distinct roles: physically “plugging” the filament tip, mediating adhesion, and driving mechanosensing. (18, 20, 24, 26, 50). We hypothesize that the PilX β-solenoid fulfills this plug function in cyanobacteria, which generally do not encode PilY1/PilC homologs. The PilX-EPS subfamily, which is encoded within the *hps* locus essential for polysaccharide secretion in *Nostoc punctiforme* and other filamentous cyanobacteria (40, 51), features particularly extensive solenoids and additional undefined domains. Given that EPS are also implicated in *Synechocystis* motility (52, 53), we propose that the PilX-EPS family acts as an adhesin that recognizes the self-laid EPS track, thereby coupling motility and surface sensing.

The distinct functions of these two machines appear to be integrated into a broader regulatory network. The inverse accumulation of Com- and EPS-related proteins implies that cells distribute competing filament types among a fixed pool of membrane complexes. This mirrors the divergent transcriptional regulation of the *pilA5-6* and *pilA9-12* operons by CRP-family regulators and second messengers (cAMP and c-di-GMP), effectively switching the shared machinery between motility and DNA uptake modes. (35, 36, 54, 55). Unexpectedly, the deletion of *pilX2* reduced EPS filaments and downregulated the expression of genes associated with the cAMP regulon. Since surface contact triggers rapid cAMP accumulation in *Synechocystis* (36), this implies that the EPS-tip complex functions as the primary apparatus for surface sensing.

Finally, the observed reduction in DnaJ3 levels in Δ*pilX2* and Δ*pilA9-12* mutant strains revealed a link between pilins and the DnaK1-DnaJ3 chaperone system that interacts with the PilB motor and is essential for motility and EPS secretion in cyanobacteria . Its suppression likely reflects a feedback mechanism that downregulates this chaperone system when the EPS pilus is compromised, further corroborating its critical link to EPS-enhanced motility.

In conclusion, this study established that *Synechocystis* assembles two genetically and morphologically distinct T4P filaments. By delineating the “Com” and “EPS” modules, we provide a robust framework for annotating these systems across the phylum. Future investigations should prioritize determining the high-resolution structures of these tip complexes and verifying their interactions with DNA or exopolysaccharides to elucidate how specific structural features dictate distinct physiological functions.

## Materials and Methods

Detailed descriptions of strain construction, primers, plasmids, bioinformatic parameters, and specific experimental protocols are provided in the SI Appendix.

### Strains and Culture Conditions

The *Synechocystis* sp. PCC 6803 substrain PCC-M (39, 44, 45, 56) and all derived mutant strains (Table S1) were cultivated on BG-11 agar plates (0.75% w/v) supplemented with 0.3% (w/v) sodium thiosulfate. For clarity, we designated the TU0763 mutant, which carries a deletion of the genes *pilA9* through *slr2019*, as Δ*pilA9-12* throughout this work. Cultures were maintained at 30°C under continuous white-light LED illumination of approx. 25 µmol photons m^-2^ s^-1^. For inducible expression, strains utilizing the *petJ* promoter were grown in CuSO_4_-free BG-11 medium, whereas strains utilizing the *nrsB* promoter were induced with 2.5 µM NiCl_2_. When required, the medium was supplemented with antibiotics at the following final concentrations: 7 µg ml^-1^ chloramphenicol, 10 µg ml^-1^ streptomycin, 50 µg ml^-1^ kanamycin, and 20 µg ml^-1^ erythromycin. *Thermosynechococcus vulcanus* NIES-2134 (57) cultures were maintained similarly but at 45°C on 1% (w/v) BG-11 agar plates and supplemented with 80 µg ml^-1^ kanamycin for selection of transformants.

### Plasmid Construction and Mutant Generation

Constructs for the generation of Δ*pilA2*, Δ*pilA4*, and Δ*pilA7-8* deletion mutants were prepared using PCR-based seamless assembly cloning. Approximately 1000 bp of the upstream and downstream regions flanking the respective genes were assembled with a kanamycin resistance cassette into a pUC19 plasmid backbone. To combine the Δ*pilA7-8* deletion with the Δ*pilA9*-*slr2019* mutant (35), the kanamycin resistance cassette in the Δ*pilA7-8* construct was replaced by an erythromycin resistance cassette. All resulting plasmids were verified by sequencing prior to transformation. For the generation of the Δ*pilA1* mutant, an existing knock-out construct (34) was used. For the generation of *Synechocystis pilA1* point mutants, a construct for the integration of a kanamycin resistance cassette between the *pilA1* and *pilA2* genes was assembled. This construct was subsequently mutated via site-directed mutagenesis to introduce the *pilA1*(T77C) mutation. In the resulting construct harboring the *pilA1*(T77C) mutation, the kanamycin selection marker was replaced by a streptomycin resistance marker. This final construct was subsequently used to create WT and minor pilin mutant strains expressing the T77C variant from the native *pilA1* locus. Constructs for the generation of Δ*tll1019*, Δ*tlr0680-82*, and Δ*tll2333-35* deletion mutants of *T. vulcanus* were prepared similar to *Synechocystis* constructs but with approximately 2500 bp of the upstream and downstream flanking regions. All resulting plasmids were verified by sequencing prior to transformation. Cyanobacterial strains were transformed with the corresponding plasmids and complete segregation of the knockouts was confirmed by colony PCR after successive rounds of antibiotic selection. For expression of minor pilin genes in *trans*, the coding sequence of *pilA8* or the sequence spanning the *pilA7-8* coding sequences was cloned into the conjugative vector pNS05 via seamless assembly. Initial attempts to express *pilA7* alone under the control of the *petJ* promoter failed, likely due to toxicity associated with its constitutive expression from the *petJ* promoter in *E. coli*. To achieve *pilA7* expression, the *petJ* promoter in pNS05 was replaced with the Ni^2+^-inducible *nrsB* promoter (58), followed by the insertion of the *pilA7* coding sequence via seamless assembly. These resulting expression plasmids were transferred into the *Synechocystis* Δ*pilA7-8* mutant strain by conjugation. Exconjugants were selected on streptomycin. For a complete list of mutant strains, plasmids, and primers used for plasmid generation, see Supplementary tables S1-S3.

### Phototaxis Assays

Phototaxis experiments were performed using non-transparent boxes with a single lateral opening for directional illumination. For *Synechocystis*, cells were restreaked onto BG-11 agar plates (0.5% agar, 0.2% glucose, 10 mM TES pH 8.0, 0.3% sodium thiosulfate), incubated for 2– 3 days under diffuse low-intensity white light, and subsequently exposed to directional white light (approx. 15 µmol photons m^−2^ s^−1^) for 7–10 days (59). For *T. vulcanus*, 5 µl of cell culture (concentrated to an OD_730_ of 30) was spotted onto BG-11 agar plates (1.0% agar, 0.3% sodium thiosulfate) and incubated under directional green LED light (approx. 120 µmol photons m^−2^ s^−1^) for 24 hours.

### Flocculation Assays

To quantify cell aggregation, cultures were diluted to an optical density (OD) at 750 _nm_ of 0.25. Aliquots of 6 ml were distributed into transparent, non-coated 6-well plates. The plates were subsequently incubated for 48 h at 30°C with 95 rpm orbital shaking and continuous illumination of approx. 30 µmol photons m^-2^ s^-1^. Aggregation was documented by imaging the chlorophyll autofluorescence using a Typhoon FLA4500 imaging system (GE Healthcare). Fluorescence was excited at 473 nm and detected at 665 nm. For each well, an aggregation score was determined by calculating the standard deviation of the autofluorescence intensity divided by the mean autofluorescence intensity.

### Transformation efficiency assay

For transformation assays, *Synechocystis* strains were grown to an OD_750nm_ of 0.8. 7.5 ml aliquots were harvested by centrifugation (2800 x g, 10 min, RT). Following the complete removal of the supernatant, cells were carefully resuspended in 100 µl BG-11 medium containing 500 ng of plasmid pNS133. Samples were subsequently incubated for 4 h at 30°C under low-intensity illumination (approx. 10-15 µmol photons m^-2^ s^-1^). The cell suspension was then plated evenly onto 50 ml 0.5% (w/v) BG-11 agar plates supplemented with 0.3% (w/v) sodium thiosulfate. After incubation for 1 day under low light, a sterile 24 mm filter disk was placed onto the center of the plate, onto which 40 µl of 10 µg ml^-1^ erythromycin was applied. Plates were incubated for two weeks, after which resistant colonies appearing in the zone of inhibition were counted. *T. vulcanus* strains were treated similarly but were cultivated at 45°C overnight before plating on 1.0% (w/v) BG-11 agar plates supplemented with 0.3% (w/v) sodium thiosulfate and 80 µg ml^-1^ kanamycin.

### Bioinformatic Workflow for Pilin Family Detection

To assemble a comprehensive dataset of putative cyanobacterial pilins, sequences were aggregated from three distinct sources. First, UniProt sequences were retrieved (accessed on 26 July 2024) based on annotation with either the Gene3D domain 3.30.700.10 (Glycoprotein, Type IV Pilin) or the InterPro entry IPR012902, which targets N-terminal methylation motifs. Second, this dataset was supplemented with sequences identified using hmmsearch (HMMER suite) to query UniProtKB. The search utilized a profile HMM built from a MAFFT (L-INS-i) alignment of *Synechocystis* PilX (*slr0226, slr0442, sll1268, slr2018*) homologs, employing iterative E-value thresholds of 1e-5 and 1e-1, respectively. Third, sequences that exhibited structural similarity (Foldseek (60) against the AlphaFold/UniRef50 database (accessed 17 March 2023); probability > 0.5) to AlphaFold2-predicted *Synechocystis* pilins were included; obvious false positives from this structural search were manually removed. For computational tractability, this combined dataset was restricted to entries from 219 cyanobacterial RefSeq genomes (accessed 20/11/2024) and five model organisms: *Synechocystis* sp. PCC 6803, *Synechococcus elongatus* PCC 7942, *Microcystis aeruginosa* NIES-843, *Nostoc* sp. PCC 7120, and *Picosynechococcus* sp. PCC 7002 (Schmelling & Bross, 2024). This filtering process yielded 985 putative pilin candidates, for 80% of which AlphaFold2 models (accessed on 7 Nov. 2025) were available and were downloaded for analysis.

To identify and annotate homologous pilin families, the topological similarity between all 794 pilin candidates was assessed using USalign (61). The Template Modeling (TM) score was calculated for all pairwise structural alignments, resulting in over 300,000 comparisons. The resulting similarity matrix was subjected to hierarchical clustering using the pvclust package in R (62), employing a correlation-based distance metric, average (UPGMA) linkage, and 10,000 bootstrap replications. Top-level clusters were considered statistically supported if they possessed an approximately unbiased (AU) p-value greater than 0.95. These supported clusters were subsequently annotated based on the enrichment of phylogenetic, family and domain annotations of their constituent sequences (Fig. S4). For further analysis cyanobacterial HpsA-like proteins (IPR049774) that exceeded the size threshold of the AlphaFold database (>1,280 aa) were assigned to the PilX_EPS_ cluster while a few small clusters lacking any recognizable pilin-like fold were excluded from the analysis.

Pairwise co-occurrence of protein families within genomic loci was assessed using a one-sided Fisher’s Exact Test to identify positive associations that significantly exceed random expectation. Significant interactions were defined as those with a Benjamini-Hochberg-adjusted p-value < 0.05 and a log2 enrichment score (observed/expected) > 1.

### Western blot analysis of surface piliation

Cultures were cultivated on BG11 agar plates (0.75% agar, 0.2% glucose, 10 mM TES pH 8, and 0.3% sodium thiosulfate). Cells were harvested by scraping, resuspended in 1 ml PBS (pH 7.4), and vortexed for exactly 1 min to shear surface pili. The cell suspensions were subsequently diluted to 1 ml aliquots of OD_750nm_ = 10 and centrifuged (30 min, 16,000 x g, RT). The resulting cell pellet, representing the PilA1 membrane fraction, was washed once with 500 µl PBS and resuspended in 100 µl 1x SDS loading buffer (250 mM Tris-HCl, pH 6.8, 40% glycerol, 4% lithium dodecyl sulfate, 2 mM EDTA, 100 mM DTT, 0.025% Bromophenol Blue). Surface pili from the 800 µl of the supernatant were precipitated by adding 100 µl 5 M NaCl and 100 µl of 30% (w/v) PEG 8000, followed by overnight incubation at 4°C. The precipitate was collected by centrifugation (1 h, 21.000 x g, 4°C), and the pellet was resuspended in 100 µl 1x SDS loading buffer. This surface pili sample was subsequently denatured for 60 min at 50°C with agitation. 7.5 µl of each sample was separated by 15% SDS-PAGE and transferred to a nitrocellulose membrane via Western blot. Membranes were blocked in 5% milk powder in TBST (20 mM Tris-HCl pH 7.5, 150 mM NaCl, 0.1% Tween-20) and PilA1 was detected using an affinity-purified anti-PilA1 antibody (1:10,000) raised in rabbits against the synthetic Ac-CGSEDTPPTPGGAND-NH2 peptide and a secondary anti-rabbit HRP-conjugate (1:20,000), followed by chemiluminescence detection. Western blot signals were quantified by densitometric analysis.

### Mass spectrometry analysis of sheared pili samples

Surface pili were sheared from *Synechocystis* strains grown for 8 days on motility plates following the isolation procedure described above, with the modification that the OD_750nm_ was adjusted to 50 prior to the removal of cells by centrifugation for 5 min. The supernatant containing the sheared pili was transferred to fresh tubes, and proteins were precipitated by the addition of 0.603 g solid ammonium sulfate followed by incubation at 4°C for 72 h. The precipitate was pelleted by centrifugation (1 h, 21.000 x g, 4°C), and the resulting protein pellet was fully resolubilized in 65 µl of 2% SDS with agitation (1,000 rpm, 10 min). Protein samples were reduced with DTT, alkylated with chloroacetamide, and subjected to automated SP3-based (63) tryptic digestion using a Hamilton robotic platform. Peptides were subsequently desalted using SDB-RPS StageTips and analyzed by liquid chromatography-mass spectrometry (LC-MS) on an Exploris 480 system coupled to an UltiMate™ 3000 RSLCnano (both Thermo Fisher Scientific) operating in data-independent acquisition (DIA) mode as described (64). Raw data processing, protein identification against a custom *Synechocystis* database, and MaxLFQ-based quantification were performed using DiaNN (v1.8.2) (65), followed by statistical analysis using the limma package. Proteins with ≥3 missing intensity values were subjected to left⍰censored imputation using the MinProb method, followed by sequential robust imputation (impSeqRob) for proteins with ≥1 remaining missing values, with all imputations performed on the valid⍰value– filtered dataset for each experimental comparison. Clustering was performed using Euclidean distance and complete linkage.

### Thiol-specific PilA1 labeling and imaging of live pili

After systematically introducing cysteine residues at various positions in the major pilin PilA1 and testing their functionality, we selected the PilA1(T77C) variant. This mutant, which confers kanamycin resistance, was confirmed to assemble into functional pili, as it retained surface-detectable PilA1, was stainable by AlexaFluor488 maleimide, and exhibited phototactic motility (Fig. S3). To combine this labeling capacity with the minor pilin deletions, the kanamycin resistance cassette in the PilA1(T77C) background was replaced with an *aadA* gene conferring streptomycin resistance. While this genetic manipulation resulted in the loss of phototactic motility, individual pilus extension and retraction events were preserved (Movies S1-7), allowing for morphological characterization. Any potential off-target effects from this cassette exchange are presumed to be comparable across all constructed mutant strains. PilA1(T77C) mutant strains were grown on motility plates for 8 d under constant illumination of approx. 20 µmol photons m^-2^ s^-1^. Cells were resuspended in 100 µl BG11, pelleted (4000 × g, 4 min, RT), and washed with sterile-filtered PBS (pH 7.4). For thiol-specific labeling, 6 µg of Alexa Fluor 488 C5 maleimide (Thermo Fisher Scientific; 1 mg/ml stock in DMSO) was added. Samples were incubated (30°C, 2 h, dark) in a thermoshaker, with 300 rpm agitation for the first hour, after which unbound dye was removed by two PBS washes. Labeled cells were resuspended in approx. 150 µl BG11 and 3 µl aliquots were spotted onto BG11 agar pads (2% agar, 0.2% glucose, 10 mM TES pH 8, and 0.3% sodium thiosulfate). Once dried, the agar spots were placed upside-down into 35 mm glass-bottom µ-Dish (ibidi) and incubated for at least 2 h at 30 °C in the dark. Images were acquired on a Nikon A1 Plus confocal microscope (Plan Apo VC 60x Oil DIC N2 objective) controlled by NIS-Elements AR software. Chlorophyll autofluorescence (Ex: 588 nm, Em: 647 nm) and Alexa Fluor staining (Ex: 493 nm, Em: 516 nm) were captured independently. Time-lapse videos were recorded for 10 s, at 0.5 frames s^-1^. For pilus quantification, the first frame was manually segmented into cells (autofluorescence), pili (Alexa Fluor), and background using ilastik (66), and a custom R script was used to remove edge-associated cells and any detected pili structures smaller than 5 pixels, as these were likely segmentation artifacts. A pairwise Wilcoxon test was used to calculate statistically significant differences between these samples. To analyze pilus dynamics, the number of pili was counted for each frame, and the total absolute change in the number of pili over all time points was calculated.

### Transmission electron microscopy

Five microliters of culture were spotted onto a formvar-coated copper grid (Plano GmbH) and incubated for 30 s. The liquid was removed using Whatman paper, and this was repeated five times. The grid was then washed with 20 µl of Milli-Q water and dried with Whatman paper. The samples were negatively stained with 2% uranyl acetate. Imaging was performed using a Hitachi HT7800 operated at 100 kV, equipped with an EMSIS Xarosa 20-MP CMOS camera.

## Acknowledgments

This work was funded by the Deutsche Forschungsgemeinschaft (DFG, German Research Foundation) – Project-ID 403222702 – SFB 1381 to AW, FD, PFA and SVA. We would like to thank the EM facility at the Faculty of Biology, University of Freiburg, for access to the TEM for generation of data. The TEM (Hitachi HT7800) was funded by the DFG grant (project number 426849454) and is operated by the University of Freiburg, Faculty of Biology, as a partner unit within the Microscopy and Image Analysis Platform (MIAP) and the Life Imaging Center (LIC), Freiburg.

## Notes

### Competing Interest Statement

The authors have declared no competing interest.

