## Supplementary Information for "Distinct core minor pilin complexes prime specialized type IV filaments in cyanobacteria"

**This PDF file includes:**

Supplementary Figures S1-S4
Supplementary Tables S1-S3
Supplementary Dataset S2

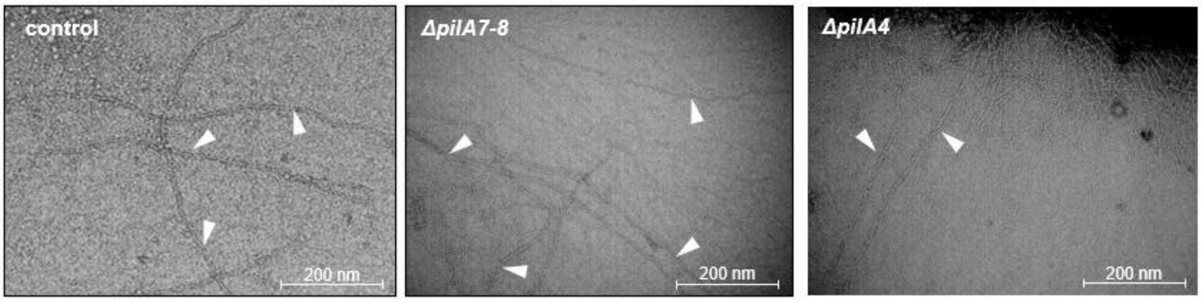


**Figure S1: TEM images of Type IV pili of *Synechocystis* pilin mutants.** Negatively stained cells (2% uranyl acetate) from a liquid culture were imaged on a Hitachi HT7800 operated at 100 kV. Representative Type IV pili are marked by white arrows. A Δ*sll1835* knockout strain, which is not related to pilus assembly, served as a control.

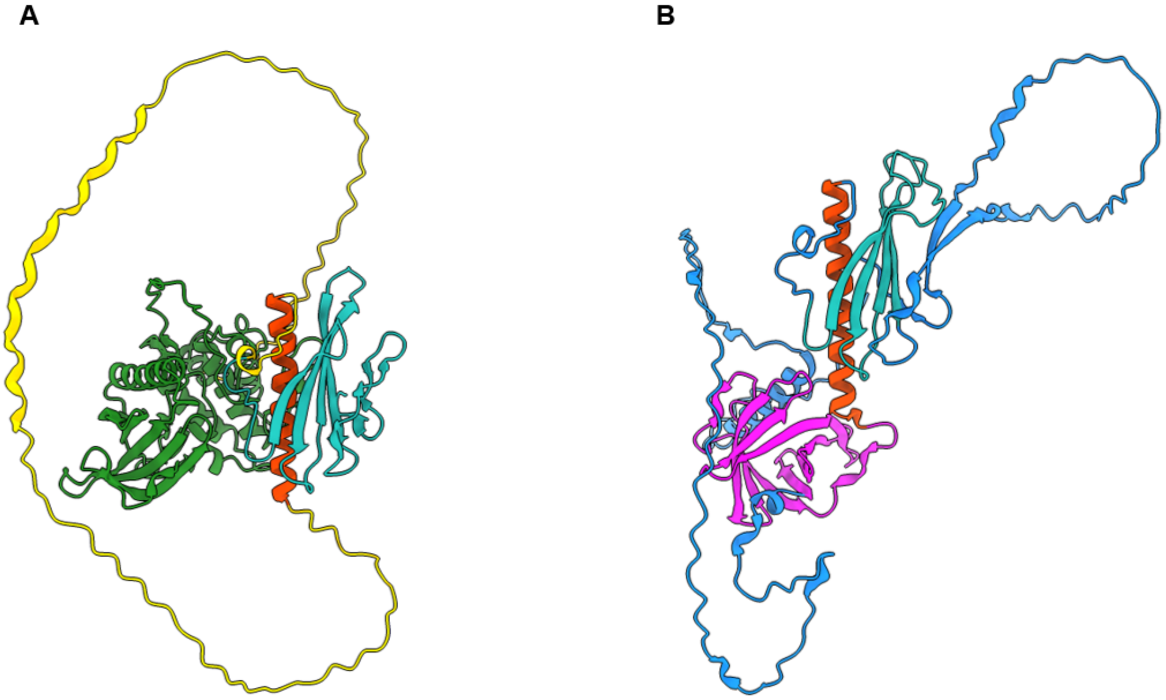


**Figure S2: Structural models of divergent pilin-like proteins.** AlphaFold2 structural predictions for two representative proteins from clades of pilin-like proteins that lack the hydrophobic α1N-helix. Despite this truncation, these proteins retain the canonical pilin core fold, characterized by the C-terminal portion of the α-helix (α1C; red) and the 4-stranded antiparallel β-sheet (turquoise). **(A)** Oscil6304_4885 from *Oscillatoria acuminata* PCC 6304 (A0A0M2Q2B0) possesses a large serine/threonine protein kinase domain (green) at the N-terminus. **(B)** PROH_03390 from *Prochlorothrix hollandica* PCC 9006 (K9TQG2) has an β-barrel domain (pink) grafted onto the pilin core.

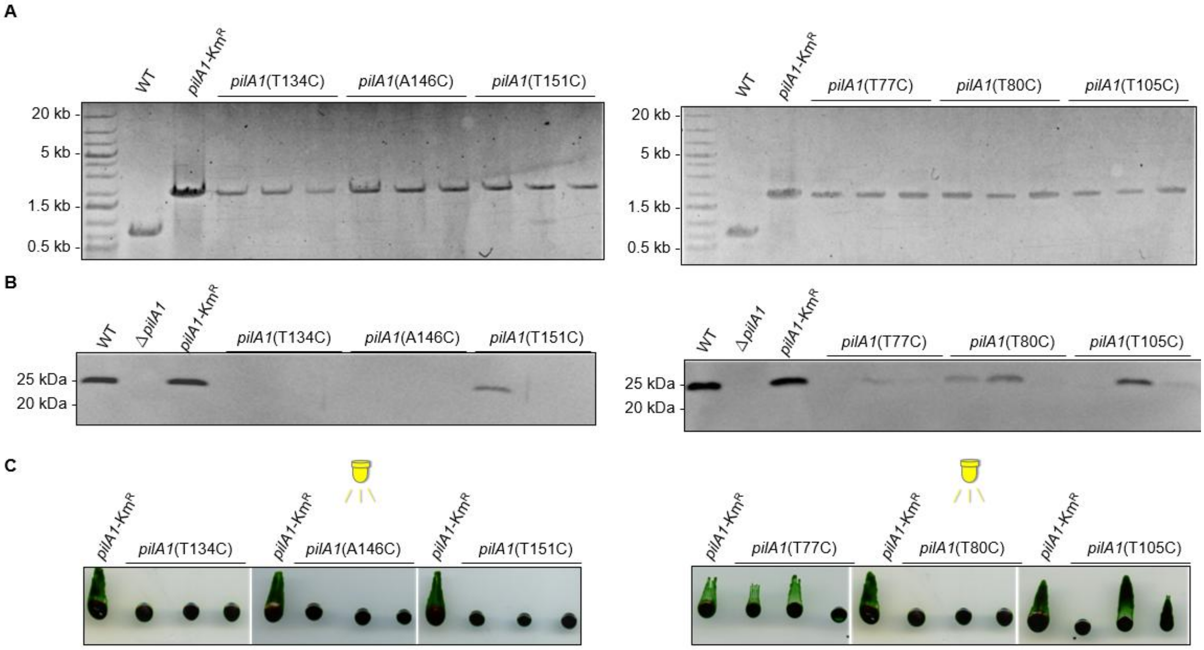


**Figure S3: Genotypic and phenotypic characterization of *Synechocystis* PilA1 cysteine variants. (A)** Genotyping of *Synechocystis* strains expressing specific PilA1 cysteine variants via colony PCR. The presence of a ~2,000 bp product indicates the integration of the pilA1:Km$^R$ cassette, distinct from the ~700 bp wild-type (WT) locus. Presence of the 700 bp band indicates incomplete segregation. **(B)** Immunoblot analysis of sheared extracellular pili fractions separated by 15% SDS-PAGE and probed with anti-PilA1 antibodies. The presence of the major pilin (~25 kDa) on the cell surface is detected in the WT, the parental PilA1-Km$^R$ strain, and mutants T77C, T80C, and T105C. Δ*pilA1* serves as a negative control. **(C)** Phototaxis assays were performed on 0.5% agar plates under directional white light (LED). Functional motility is observed for T77C, T80C, and T105C mutants, whereas T134C, A146C and T151C variants exhibit a non-motile phenotype similar to the Δ*pilA1* mutant.

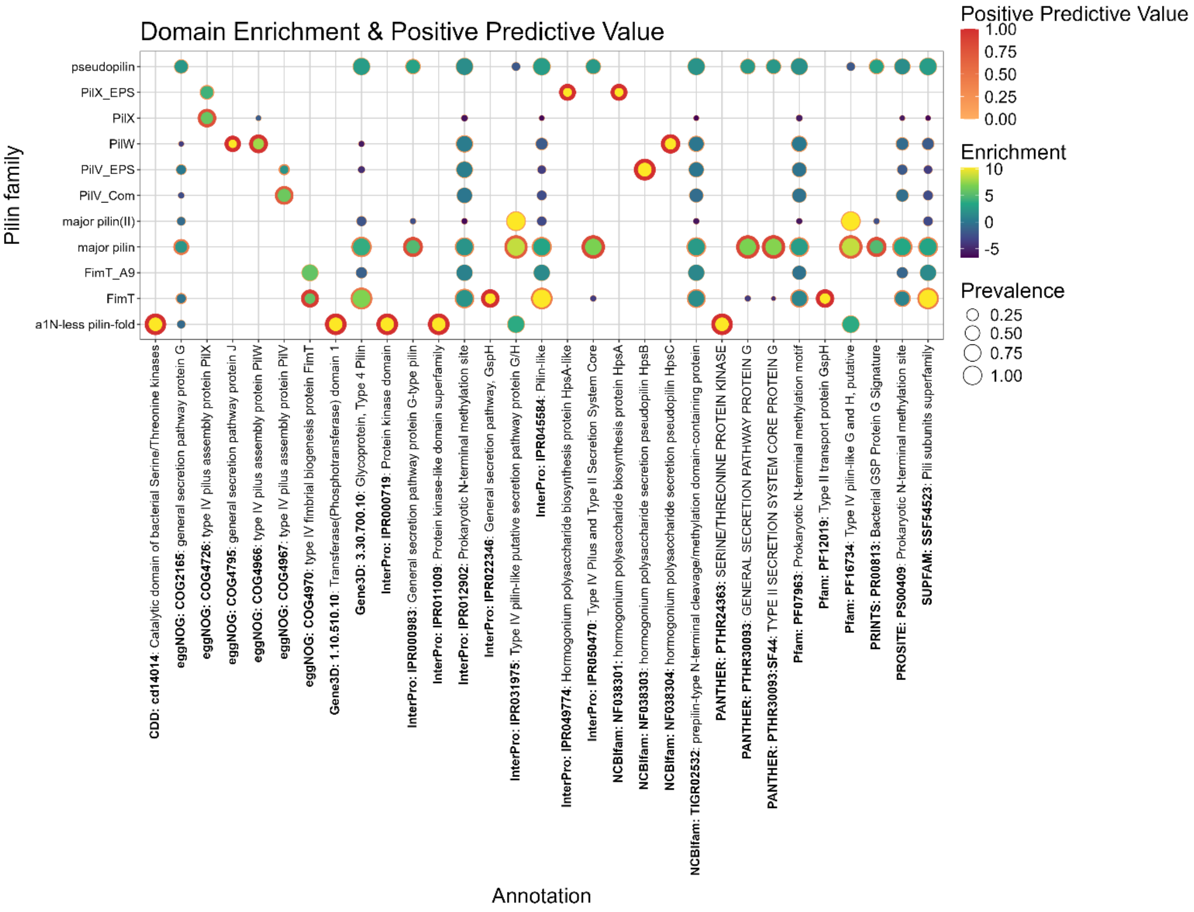


**Figure S4: Identification of signature domains across pilin families**. The Bubble plot displays the enrichment and specificity of protein annotations (InterPro, eggNOG, Pfam, etc.) associated with specific pilin families. The x-axis displays protein annotations that ranked among the top five significant annotations for at least one pilin family defined by the cluster analysis (y-axis). Fill color denotes the enrichment strength (Log2 Odds Ratio) relative to the dataset background derived from a Fisher's exact test; Bubble size represents prevalence, defined as the proportion of proteins within a locus possessing the annotation. Border thickness and color indicate the positive predictive value (PPV), the probability that a protein belongs to the locus given the presence of the annotation.

**Table S1: Strains used in this study**

| Strain | Parent | Relevant Genotype | Reference/Plasmid |
|---|---|---|---|
| Wild Types & existing strains | | | |
| *Synechocystis* PCC-M | N/A | Wild type | (1) |
| Δ*pilA5-6* | WT | TU2300::Cm | (2) |
| Δ*pilA9-12* | WT | TU763::Km | (2) |
| *Thermosynechococcus vulcanus* NIES-2134 | N/A | Wild type | (3) |
| *Synechocystis* strains created for this study | | | |
| Δ*pilA1* | WT | *pilA1*::Cm$^R$ | pGEM-Teasy-ΔpilA1-CamR |
| Δ*pilA2* | WT | *pilA2*::Km$^R$ | pNS112 |
| Δ*pilA4* | WT | *pilA4*::Km$^R$ | pNS061 |
| Δ*pilA7-8* | WT | *pilA7-8*::Km$^R$ | pNS062 |
| Δ*pilA7-8*/*pilA7*$^+$ | Δ*pilA7-8* | *pilA7-8*::Km$^R$ / [P*rnsB*::*pilA7*, Sm$^R$] | pNS142 |
| Δ*pilA7-8*/*pilA8*$^+$ | Δ*pilA7-8* | *pilA7-8*::Km$^R$ / [P*petJ*::*pilA8*, Sm$^R$] | pNS132 |
| Δ*pilA7-8*/*pilA7-8*$^+$ | Δ*pilA7-8* | *pilA7-8*::Km$^R$ / [P*petJ*::*pilA7-8*, Sm$^R$] | pNS131 |
| Δ*pilA9-12*/*pilA5-6* | Δ*pilA9-12* | TU763::Km$^R$, TU2300::Cm$^R$ | pJET-ΔTU2300-CmR-rev |
| Δ*pilA9-12*/*pilA7-8* | Δ*pilA9-12* | TU763::Km$^R$, *pilA7-8*::Em$^R$ | pNS133 |
| *pilA1*-Km$^R$ | WT | *pilA1*:Km$^R$ | pUC-*pilA1*-KmR |
| *pilA1*(T77C) | WT | *pilA1*(T77C)-Km$^R$ | pNS074 |
| *pilA1*(T80C) | WT | *pilA1*(T80C)-Km$^R$ | pNS075 |
| *pilA1*(T105C) | WT | *pilA1*(T105C)-Km$^R$ | pNS077 |
| *pilA1*(T134C) | WT | *pilA1*(T134C)-Km$^R$ | pNS087 |
| *pilA1*(A146C) | WT | *pilA1*(A146C)-Km$^R$ | pNS076 |
| *pilA1*(T151C) | WT | *pilA1*(T151C)-Km$^R$ | pNS078 |
| WT$^{T77C}$ | WT | *pilA1*(T77C)-Sm$^R$ | pNS121 |
| Δ*pilA5-6*$^{T77C}$ | WT$^{T77C}$ | TU2300::Cm$^R$, *pilA1*(T77C)-Sm$^R$ | pJET-ΔTU2300-CmR-rev |
| Δ*pilA7-8*$^{T77C}$ | WT$^{T77C}$ | Δ*pilA7-8*::Km$^R$, *pilA1*(T77C)-Sm$^R$ | pNS133 |
| Δ*pilA9-12*$^{T77C}$ | Δ*pilA9-12* | TU763::Km$^R$, *pilA1*(T77C)-Sm$^R$ | pNS121 |
| Δ*pilX1*$^{T77C}$ | WT$^{T77C}$ | *slr0226*::Gm$^R$, *pilA1*(T77C)-Sm$^R$ | pNS149 |
| Δ*pilX2*$^{T77C}$ | WT$^{T77C}$ | *slr0442*::Km$^R$, *pilA1*(T77C)-Sm$^R$ | pJET-Δslr0442 |
| Δ*pilX3*$^{T77C}$ | WT$^{T77C}$ | *sll1268*::Em$^R$, *pilA1*(T77C)-Sm$^R$ | pNS150 |
| Δ*tll1019* | WT | Δ*tll1019::Km$^R$* | pUC19-Δtll1019_Kmv2 |
| Δ*tlr0680-82* | WT | Δ*tlr0680-82::Km$^R$* | pUC19-Δtlr0680-82_Kmv2 |
| Δ*tll2333-35* | WT | Δ*tll2333-35::Km$^R$* | pUC19-Δtll2333-35_Kmv2 |

**Table S2: Plasmids used in this study**

| Plasmid | Description / Function | Source / Reference |
|---|---|---|
| Plasmids (This Study) | | |
| pNS05 | RSF1010-Based conjugative expression vector with P*petJ* | This study |
| pNS141 | pNS05 derivative with P*rnsB* promoter | This study |
| pNS142 | P*rnsB*-driven expression of *pilA7* | This study |
| pNS131 | P*petJ*-driven expression of *pilA7-pilA8* | This study |
| pNS132 | P*petJ*-driven expression of *pilA8* | This study |
| pNS062 | Knockout construct for *pilA7-pilA8* (*pilA7-8*::Km$^R$) | This study |
| pNS133 | Knockout construct for *pilA7-pilA8* (*pilA7-8*::Em$^R$) | This study |
| pNS112 | Knockout construct for *pilA2* (*pilA2*::Km$^R$) | This study |

| | | |
|---|---|---|
| pNS061 | Knockout construct for *pilA4* (*pilA4*::Km$^R$) | This study |
| pUC-pilA1-KmR | Construct for Km$^R$ integration between *pilA1* and *pilA2* | This study |
| pNS074 | pUC-pilA1-KmR derivative with *pilA1(T77C)* | This study |
| pNS075 | PUC-pilA1-KmR derivative with *pilA1(T80C)* | This study |
| pNS077 | PUC-pilA1-KmR derivative with *pilA1(T105C)* | This study |
| pNS087 | PUC-pilA1-KmR derivative with *pilA1(T134C)* | This study |
| pNS076 | PUC-pilA1-KmR derivative with *pilA1(A146C)* | This study |
| pNS078 | PUC-pilA1-KmR derivative with *pilA1(T151C)* | This study |
| pNS121 | pNS074 derivative where Km$^R$ is replaced by Sm$^R$ | This study |
| pNS149 | pJET-Δ*slr0226* derivative where Gm$^R$ replaces Km$^R$ | This study |
| pNS150 | pJET-Δ*sll1268* derivative where Km$^R$ is replaced by Em$^R$ | This study |
| pUC19-Δ*tll1019*_Kmv2 | Knockout construct for NIES2134_108990 | This study |
| pUC19-Δ*tlr0680*-82_Kmv2 | Knockout construct for NIES2134-105520 | This study |
| pUC19-Δ*tll2333*-35_Kmv2 | Knockout construct for NIES2134-105520 | This study |
| Source Plasmids | | |
| pUR | Source of streptomycin resistance (Sm$^R$) cassette | (1) |
| pUC-Δ*pixGH* | Source of gentamycin resistance (Gm$^R$) cassette | (4) |
| pUC4K | Source of kanamycin resistance (Km$^R$) cassette | Amersham |
| pUC19 | Amp$^R$ backbone for cloning knockout constructs | (5) |
| pGEM-Teasy-ΔpilA1-CamR | Construct for *pilA1* knockout (*pilA1*::Cm$^R$) | (6) |
| pJET-Δ*slr0226* | Knockout construct for *slr0226* (*slr0226*::Km$^R$) | (7) |
| pJET-Δ*sll1268* | Knockout construct for *sll1268* (*sll1268*::Km$^R$) | (7) |
| pJET-Δ*slr0442* | Knockout construct for *slr0442* (*slr0442*::Km$^R$) | (7) |
| pJET-ΔTU2300-CmR-rev | Knockout construct for TU2300 (*pilA5-6*::Cm$^R$) | (2) |

**Table S3: Oligonucleotides used in this study**

| Primer Name | Sequence (5' to 3') | Template/Purpose |
|---|---|---|
| **pNS141 Construction** | | |
| NS310 | TCATGATCTTTATAATCCATACCACCTCAAATTGGGAATT | *rnsB* promoter |
| NS311 | CGGTCAAGGTTCTGGACCAGTTCCACCAGCAAAATTCGCA | *rnsB* promoter |
| NS312 | AATTCCCAATTTGAGGTGGTATGGATTATAAAGATCATGATGGCG | pNS05 |
| NS313 | TGCGAAATTTTGCTGGTGGAACTGGTCCAGAACCTTGACCG | pNS05 |
| **pNS142 Construction** | | |
| NS314 | AATTCCCAATTTGAGGTGGTATGGCTTACAAGAATGAACTG | *pilA7* |
| NS315 | GGCAACCGAGCGTTGGATCCCTAATAACCAGTTCTGCATTGC | *pilA7* |
| NS316 | AATGCAGAACTGGTTATTAGGGATCCAACGCTCGGTTGCC | pNS142 |
| NS317 | AGTTCATTCTTGTAAGCCATACCACCTCAAATTGGGAATTTGTC | pNS142 |
| **pNS131 Construction** | | |
| NS291 | AGTTCATTCTTGTAAGCCATATGTTCTCCTTCAAGGATAAAG | pNS05 |
| NS292 | GTACAATTAATGCAGTCTTTAATAAGGATCCAACGCTCG | pNS05 |
| NS293 | CGAGCGTTGGATCCTTATTAAAGACTGCATTAATTGTACAAC | *pilA7-pilA8* |
| NS294 | TATCCTTGAAAGGAGAACATATGGCTTACAAGAATGAACTG | *pilA7-pilA8* |
| **pNS132 Construction** | | |
| NS295 | GTACAATTAATGCAGTCTTTAATAAGGATCCAACGCTCG | pNS05 |
| NS296 | AAGTACCATAGAATTTTCACATGTTCTCCTTCAAGGATAAAG | pNS05 |
| NS297 | TATCCTTGAAAGGAGAACATGTGAAAATTCTATGGTTACTAATGAC | *pilA8* |
| NS298 | CGAGCGTTGGATCCTTATTAAAGACTGCATTAATTGTACAAC | *pilA8* |
| **pNS062 Construction** | | |
| NS050 | GTGCGGGCCTCTTCGCTATTTTTGAGCAGGTTGAAAAACA | *pilA8* flank |

| | | |
|---|---|---|
| NS051 | AGATTTTGAGACACAACGTGAGCAAAAATGGTAGCTTAAAATTAT | *pilA8* flank |
| NS052 | TTTAAGCTACCATTTTTGCTCACGTTGTGTCTCAAAATCT | Km$^R$ cassette |
| NS053 | CTTGAACAAGACTTTGCATTGATCCTTCAACTCAGCAAAA | Km$^R$ cassette |
| NS054 | TTTTGCTGAGTTGAAGGATCAATGCAAAGTCTTGTTCAAGATC | *pilA7* flank |
| NS055 | GAGTTAGCTCACTCATTAGGAGTTCGGGCATTAGCTGCAA | *pilA7* flank |
| NS056 | TTGCAGCTAATGCCCGAACTCCTAATGAGTGAGCTAACTCACATT | pUC19 backbone |
| NS057 | TGTTTTTCAACCTGCTCAAAAATAGCGAAGAGGCCCGCAC | pUC19 backbone |
| **pNS133 Construction** | | |
| NS299 | AATTAGTATAATTATAGCACGAGCAAAAATGGTAGCTTAAAATTATCT | pNS062 |
| NS300 | GTTAAGGGATGCAGGTCGACAATGCAAAGTCTTGTTCAAGA | pNS062 |
| NS301 | CTTGAACAAGACTTTGCATTGTCGACGTGCATCCCTTAAC | Em$^R$ cassette |
| NS302 | TTTAAGCTACCATTTTTGCTCGTGCTATAATTATACTAATTTTATAAGGA | Em$^R$ cassette |
| **pNS112 Construction** | | |
| NS174 | AAAAAACGACGTAGATCCATCACCGAAACGCGCGAGACGA | pUC19 backbone |
| NS175 | GCCCCAATAAACATATTCTTCACTGACTCGCTGCGCTCGG | pUC19 backbone |
| NS176 | TCGTCTCGCGCGTTTCGGTGATGGATCTACGTCGTTTTTTC | *pilA2* flank |
| NS177 | AGATTTTGAGACACAACGTGAACAATAGTGTGAAAATATTAACCCA | *pilA2* flank |
| NS178 | AATATTTTCACACTATTGTTCACGTTGTGTCTCAAAATCT | Km$^R$ cassette |
| NS179 | AATTCAGCTTAAATACCATATTAGAAAAACTCATCGAGCATC | Km$^R$ cassette |
| NS180 | TGCTCGATGAGTTTTTCTAATATGGTATTTAAGCTGAATTTACACG | *pilA2* flank |
| NS181 | CCGAGCGCAGCGAGTCAGTGAAGAATATGTTTATTGGGGCAC | *pilA2* flank |
| **pNS061 Construction** | | |
| NS042 | GTGCGGGCCTCTTCGCTATTAAGTTGCCGTGGTGAATAGT | *pilA4* flank |
| NS043 | AGATTTTGAGACACAACGTGCTCGGTCTTCATAGCTGTGC | *pilA4* flank |
| NS044 | GCACAGCTATGAAGACCGAGCACGTTGTGTCTCAAAATCT | Km$^R$ cassette |
| NS045 | AGTAATCCATCCCTAGAGCCGATCCTTCAACTCAGCAAAA | Km$^R$ cassette |
| NS046 | TTTTGCTGAGTTGAAGGATCGGCTCTAGGGATGGATTACT | *pilA4* flank |
| NS047 | GAGTTAGCTCACTCATTAGGTAAACATTTAACTATTCGACAAAATATTG | *pilA4* flank |
| NS048 | GTCGAATAGTTAAATGTTTACCTAATGAGTGAGCTAACTCACATT | pUC19 backbone |
| NS049 | ACTATTCACCACGGCAACTTAATAGCGAAGAGGCCCGCAC | pUC19 backbone |
| **pUC-*pilA1*-KmR Construction** | | |
| pUC-HR1-fw2 | AAAACGACGGCCAGTGAATTCTGAGTTCCATGTGC | *pilA1* flank |
| pUC-HR1-rv2 | AAAACGACGGCCAGTGAATTCTGAGTTCCATGTG | pUC19 backbone |
| HR1-Km-fw2 | AGTAATTAAATAGGACGGGAAAGCCACGTTGTG | Km$^R$ cassette |
| HR1-Km-rv2 | ACAACGTGGCTTTCCCGTCCTATTTAATTACTTCAGC | *pilA1* flank |
| Km-HR2-fw2 | TCACGAGGCAGACCTCCCTATTATGTTTTGAGTG | *pilA2* flank |
| Km-HR2-rv2 | CTCAAAACATAATAGGGAGGTCTGCCTCGTGAAG | Km$^R$ cassette |
| HR2-pUC-fw2 | TTAGTATCCGAGAAAGGATCCTCTAGAGTCGACC | pUC19 backbone |
| HR2-pUC-rv2 | GGTCGACTCTAGAGGATCCTTTCTCGGATACTAA | *pilA2* flank |
| **Site-directed mutagenesis** | | |
| NS084 | TGAGAAGGGTTGTTTTGCAACCGATACG | *pilA1(T77C)* |
| NS085 | GTAAAATAACCTTGTTGG | *pilA1(T77C)* |
| NS086 | ACGTTTGCATGTGATACGGAAACGCTTG | *pilA1(T80C)* |
| NS087 | ACCCTTCTCAGTAAAATAACC | *pilA1(T80C)* |
| NS088 | GCTGACAACTGTGAAGCTATCCAAGAC | *pilA1(T105C)* |
| NS089 | AGTGTTAACTGCAAAAC | *pilA1(T105C)* |
| NS090 | CGATTCAGGCTGTCGCGCCTTCAGTAC | *pilA1(T134C)* |
| NS091 | TAAAACGTTCCACCTGAC | *pilA1(T134C)* |
| NS092 | TCGGGCTGAATGTGGTAGTGAAGATACTC | *pilA1(A146C)* |
| NS093 | CAAACAACAGTACTGAAGG | *pilA1(A146C)* |
| NS094 | TAGTGAAGATTGTCCCCCAACCCCAGG | *pilA1(T151C)* |

| | | |
|---|---|---|
| NS095 | CCTGCTTCAGCCCGACAAAC | *pilA1(T151C)* |

**pNS121 Construction**

| | | |
|---|---|---|
| NS210 | CGGTCAAGGTTCTGGACCAGGTCCTATTTAATTACTTCAGCACC | pUC-*pilA1*-KmR |
| NS211 | TTTTGGCGCGCCGCTAGCATCCTATTATGTTTTGAGTGGTGC | pUC-*pilA1*-KmR |
| NS212 | ACCACTCAAAACATAATAGGATGCTAGCGGCGCGCAAAA | $Sm^R$ cassette |
| NS213 | CTGAAGTAATTAAATAGGACCTGGICCAGAACCTTGACCGA | $Sm^R$ cassette |

**pNS149 Construction**

| | | |
|---|---|---|
| NS334 | CCAATTCGAGCTCGGTACCCTTAGTTGTTTTTCTAGAAGTTTTCTTGCTACC | pJET-Δ*slr0226* |
| NS335 | TCAATTCGAGCTCGGTACCCAGGGAGCGACTTCAGCCCAC | pJET-Δ*slr0226* |
| NS336 | GTGGGCTGAAGTCGCTCCCTGGGTACCGAGCTCGAATTGA | $Gm^R$ cassette |
| NS337 | ACTTCTAGAAAAACAACTAAGGGTACCGAGCTCGAATTGG | $Gm^R$ cassette |

**pNS150 Construction**

| | | |
|---|---|---|
| VR5 | ATTTCTTTTTTCGTCGACTTAAAAGCAACCAGCAGG | pJET-Δ*sll1268* |
| VR6 | CCTGCTGGTTGCTTTTAAGTCGACGAAAAAAGAAAT | $Em^R$ cassette |
| VR7 | AAGGATAAAACTGAACTTACTTACTTATTAAATAAT | $Em^R$ cassette |
| VR8 | ATTATTTAATAAGTAAGTAAGTTCAGTTTTATCCTT | pJET-Δ*sll1268* |

**pUC19-Δtvtll1019_Kmv2 Construction**

| | |
|---|---|
| pUC19-45R_tvtll1019 | AATTTGGGCAATGGGATACCGAGCTCGAATTCAC |
| pUC19-46F_tvtll1019 | TGCAAACGCTGTGAAATGCAAGCTTGGCGTAATC |
| tvtll1019-1F_pUC19 | GAATTCGAGCTCGGTATCCCATTGCCCAAATTCG |
| tvtll1019-2R_Kmv2 | CCTGAGTGCTTGCGGCTCGATAGAACCAAGGCAG |
| Km-23F_tvtll1019 | GCCTTGGTTCTATCGAGCCGCAAGCACTCAGG |
| Km-24R_tvtll1019 | TATGTACGTGGCGTAGCCTTTCATAGAAGGCGGC |
| tvtll1019-3F_Kmv2 | CGCCTTCTATGAAAGGCTACGCCACGTACATAAC |
| tvtll1019-4R_pUC19 | TTACGCCAAGCTTGCATTTCACAGCGTTTGCAGC |

**pUC19-Δtvtlr0680-82_Kmv2 Construction**

| | |
|---|---|
| pUC19-47R_tvtlr0680 | AGTCACACCATCAAAGTACCGAGCTCGAATTCAC |
| pUC19-48F_tvtlr0680 | TCCTTGAGAAAGGTGGTGCAAGCTTGGCGTAATC |
| tvtlr0680-1F_pUC19 | GAATTCGAGCTCGGTACTTTGATGGTGTGACTCC |
| tvtlr0680-2R_Kmv2 | CCTGAGTGCTTGCGGCTCTCCGTCAAGGTAAAGC |
| Km-25F_tvtlr0680 | TTTACCTTGACGGAGAGCCGCAAGCACTCAGG |
| Km-26R_tvtlr0680 | AGATACTCCAGTGGTTCCTTTCATAGAAGGCGGC |
| tvtlr0680-3F_Kmv2 | CGCCTTCTATGAAAGGAACCACTGGAGTATCTGC |
| tvtlr0680-4R_pUC19 | TTACGCCAAGCTTGCACCACCTTTCTCAAGGATC |

**pUC19-Δtvtll2333-35_Kmv2 Construction**

| | |
|---|---|
| pUC19-49R_tvtll2333 | AGGGCATTGTGTTTGTTACCGAGCTCGAATTCAC |
| pUC19-50F_tvtll2333 | AGAACTGGAACGCTATTGCAAGCTTGGCGTAATC |
| tvtll2333-1F_pUC19 | GAATTCGAGCTCGGTAACAAACACAATGCCCTCC |
| tvtll2333-2R_Kmv2 | CCTGAGTGCTTGCGGCGGGCTTGTATCCCATTTC |
| Km-27F_tvtll2333 | AATGGGATACAAGCCCGCCGCAAGCACTCAGG |
| Km-28R_tvtll2333 | AAACCACACCCGTTGACCTTTCATAGAAGGCGGC |
| tvtll2333-3F_Kmv2 | CGCCTTCTATGAAAGGTCAACGGGTGTGGTTTAC |
| tvtll2333-4R_pUC19 | TTACGCCAAGCTTGCAATAGCGTTCCAGTTCTCC |
